# Floral Reversion based micropropagation of Day-Neutral *Cannabis sativa* L

**DOI:** 10.1101/2024.09.19.613882

**Authors:** Darya Sadat Tabatabaei, Rambod Abiri, A.M.P. Jones

## Abstract

Micropropagation systems have been developed for photoperiod sensitive cultivars of cannabis, but there are currently no published methods for day-neutral genotypes. Most established cannabis micropropagation systems rely on shoot proliferation using modal explants that need them to be maintained in vegetative growth, which is not possible for day-neutral genotypes. Floral reversion, the process by which plants revert to vegetative growth from the generative stage, has been demonstrated as an alternative and potentially more efficient approach to the micropropagation of photoperiodic cannabis. The current study investigated if this process could be adapted to facilitate the micropropagation of day-neutral genotypes and overcome existing barriers. During the process, various factors such as photoperiod and light intensity may influence the efficiency of floral reversion. To evaluate this approach in day-neutral cultivars, various photoperiods were compared to assess the impact on reversion rate and subsequent growth. Excised florets obtained from *in vitro* day-neutral *C. sativa* cv. “Blue Auto Mazar x auto Blueberry” seedlings were subjected to five photoperiods including 12.0, 16.0, 18.0, 20.0, and 24.0 hours of light per day for four weeks. Reversion rates and time, shoot length, shoot number, and node number were measured. Statistical analyses revealed significant differences (p-values < 0.05) in terms of reversion time among photoperiod treatments. The highest reversion rate happened under the 16.0 hr photoperiod with 72% success. The highest shoot lengths were observed under 20.0 hr of photoperiods with a mean of 10.1 mm and the lowest shoot length belonged to 12.0 hr of photoperiods with a mean of 5.6 mm, respectively. Reverted shoots developed vegetatively for some time before initiating new flowers. However, during this period the shoots were successfully rooted in vitro and then acclimated to the growth chamber where they completed their life cycle, including seed production. This process was also successfully achieved from a 2-year old culture of *C. sativa* cv. “Blue Auto Mazar”, demonstrating that it is feasible to use this approach for long term culture maintenance and multiplication. This study demonstrates that floral reversion can successfully be used to clonally propagate day-neutral cannabis plants and serves as a basis for developing large-scale clonal propagation and breeding strategies.

## Introduction

*Cannabis sativa L*. is in the Cannabaceae family., cultivated for therapeutic and recreational purposes (Anna K. Jäger a, 1995; Bonini et al., 2018; Kalant, 1972). Different cultures have utilized this versatile plant for thousands of years, not only for its medicinal properties, but also for its nutritional value, sustainable industrial applications, and potential in renewable energy sources. Over the past decade, more countries have legalized cannabis fand it has since developed into a multi-billion dollar industry (Monthony et al., 2021).

Most Cannabis cultivars used for medicinal or recreational applications are dioecious and short-day plants, but day-neutral genotypes are becoming more common (Hammond & Mahlberg, 1977). Although male plants have fertilization purposes, the primary psychoactive and pharmaceutical substances are sequestered into glandular trichomes that are predominantly found on flowering female plants (Livingston et al., 2020). Previous studies of photoperiodic *C. sativa* in both in-vitro and ex-vitro conditions indicate that flowering can be induced under short photoperiod 12/12 hours light/dark (Ahrens et al., 2023; Babaei et al., 2022; Moher et al., 2021). Solitary flowers with stigma developing at the leaf axis and three pairs of florets with stigmas at the apical shoot indicate that the plant has shifted to the flowering stage (Spitzer-Rimon et al., 2022). Matured female plants form a highly branched compound raceme structure in the shoot apex with intense branching (Spitzer-Rimon et al., 2019; Hesami et al., 2023).

While cannabis is a prolific seed producer and feminized seeds can be used to generate all female populations, most producers rely on clonal propagation to avoid the heterogeneity of seed populations (Hesami et al., 2021). Conventional clonal propagation methods rely on stem cuttings taken from mother plants maintained under long days in the vegetative stage (Caplan et al., 2018) Alternatively, cannabis can be clonally propagated using micropropagation techniques, promising more efficient and reliable methods for cannabis breeders to produce high-quality, insect/disease-free propagules (Adhikary et al., 2021). Most published micropropagation systems for cannabis rely on shoot proliferation from nodal explants. Nodal micropropagation protocols rely on vegetative meristem that exists in the apical and axillary nodes with each plant producing a limited number, typically 3-4 nodes/plantlet (Monthony et al., 2021). As mentioned, most cannabis cultivars are photoperiod-sensitive, allowing them to be maintained in the vegetative stage of growth indefinitely. This allows repeated cycles of shoot proliferation during stage 2 of micropropagation (multiplication) to obtain an exponential increases in the number of plants propagated from limited material.

More recently, day-neutral (aka auto-flowerers), have gained more popularity as they can quickly transition into the flowering stage regardless of photoperiod and open new possibilities for cannabis producers. Unlike photoperiod-sensitive cultivars, which often fail to mature in time for harvest in northern climates, day-neutral cultivars can flower during the summer months to expand the geographical range for outdoor cannabis cultivation. In southern regions with longer growing seasons, the early planting of day-neutral cultivars in spring can facilitate an early harvest, allowing for the potential of a second crop and thereby enhancing production efficiency. Furthermore, sequential planting of day-neutral cultivars enables staggered harvests, effectively distributing labour and infrastructure demands over time, which further optimizes production efficiency. While some of these benefits can also be achieved through methods such as light deprivation and other controlled interventions, day-neutral cultivars offer a more streamlined and potentially cost-effective approach. However, this trait also introduces several challenges, as they are unable to remain in a vegetative condition, making them difficult to propagate clonally and maintain in tissue culture. This challenge has significant implications as most seedling populations are highly heterogeneous, limiting the application of seed based cultivation. Additionally, because the final product is unfertilized female flowers, by the time a plant is phenotyped, it is too late to use it to make F1 crosses. In photoperiodic cannabis, this is overcome by taking a clone prior to flowering that can be used for future crosses, but this is not currently possible with day-neutral genotypes (Monthony et al., 2021; Piunno et al., 2019).

Floral reversion, a process in which plants revert from flowering to the vegetative growth, has been introduced to overcome *in vitro* challenges such as poor multiplication rates, severe contamination, and a lack of plant material that might impede traditional multiplication methods in a variety of species (Ahmad et al., 2011; Poluboyarova et al., 2014; Supaibulwatana & Mii, 1997; Wyka et al., 2006). The structure of the cannabis inflorescence makes it a great alternative starting material compared to nodal explants due to the high density of meristems produced within the inflorescence that could potentially be used to generate plants (Spitzer-Rimon et al., 2019). The first floral reversion protocol developed for photoperiod-sensitive cannabis showed the possibility of floral reversion using greenhouse-grown florets. This approach successfully produced true-to-type plants and may serve as an alternative to maintaining clonal copies during the breeding process or in cases where the clone is lost (Piunno et al., 2019). Subsequent research demonstrated that cannabis produces flowers *in vitro* under short day conditions (Moher et al., 2021) and that these florets can be used to generate shoots through floral reversion (Monthony, et al., 2021). On average, *in vitro* plants were found to produce approximately 20 florets per plant, compared to 3-4 nodes in vegetative plants, suggesting that the multiplication rate using floral explants may be much greater.

All previous floral reversion studies have focused exclusively on photoperiod-sensitive cultivars and no prior research has explored this process in day-neutral cultivars of cannabis or examined the impact of varying photoperiods on floral reversion. This study aimed to build on previous floral reversion protocols and assess the potential of using this approach for the micropropagation of day-neutral genotypes. To the best of our knowledge, this represents the first report of day-neutral cannabis micropropagation and will serve as a basis to improve breeding and propagation methods.

## Material and Method

### Plant material

Seeds of *Cannabis sativa* cv. “Blue Auto Mazer x Auto Blueberry” (Dutch Passion, The Netherlands), were used in this study. In addition, two-year-old cultures of *C. sativa* c. “Blue Auto Mazar” were maintained *in vitro* using floral reversion, recovered and acclimatized using the developed method.

### Seed germination, media composition and culture condition

*In vitro* seed germination was conducted as described by Hesami et al, (2021). In brief, the seeds were soaked in distilled water for 48 hours in the dark. Afterward, the seeds were transferred into 10 % (v/v) commercial bleach (Purox, Kik Holdco Company Inc, Ontario) and agitated for 15 min, followed by three 5 min rinses in autoclaved distilled water under aseptic conditions. Next, seeds were placed individually into GA7 vessels (Magenta LLC, Chicago, IL) containing 40 ml of half strength Murashige and Skoog medium Van der Salm modification (M5642, PhytoTech Labs, Kansas, USA) supplemented with 23 g/L sucrose and 6 g/L agar (Thermo-Fisher Scientific, Waltham, MA). After 4 weeks, each plant was sub-cultured individually into GA7 vessels with 40 ml of basal DKW media, comprised of 5.32 g/L Driver and Kuniyuki Walnut (DKW) basal salt supplemented with vitamins (D2470, PhytoTech Labs, Kansas, USA) 30 g/L sucrose, 1 mL/L plant preservative mixture (PPM) and 6 g/L of agar.

### *In vitro* culture condition

Single day-neutral plants in Magenta GA7 vessel were maintained under 16/8 h (light/dark) photoperiod with LEDs providing a photosynthetic photon flux density (PPFD) of 30 ± 2 μmol·m–2·s–1 at 25 °C. Florets with white stigma emerged from the apical shoots after they were transferred into DKW media and were used as the starting material for this experiment.

### Floral Reversion media composition and culture condition

Floral explants were carefully dissected as pairs using scalpels and forceps, and cultured in 100 × 20 mm Petri dishes (VWR International, Mississauga, Canada). Each Petri plate contained approximately 25 mL of semi-solid medium consisting of 5.32 g/L DKW basal salt with vitamins, 30 g/L sucrose, 1 mL/L plant preservative mixture (PPM) and 6 g/L of agar with 1.0 μM meta-topolin hormone (added via filter sterilization) as described by (Monthony et al., 2021). Each petri dish contained five floral explants with each pair counting as one explant (total 125 subsamples) and five replicates (Petri dishes) per photoperiod treatment. Five photoperiod treatments of 12/12, 16/8, 18/6, 20/4, and 24/0 hours (light/dark), with light emitting diode (LED), provided a photosynthetic photon flux density (PPFD) of approximately 50 ± 2 mmol·m– 2·s^−1^ at 25 °C. PPFD and light spectrum were measured and calibrated with a Li-Cor LI-180 (LICOR, Lincoln, NE) at plant height (Fig. 1s).

**Fig 1.**
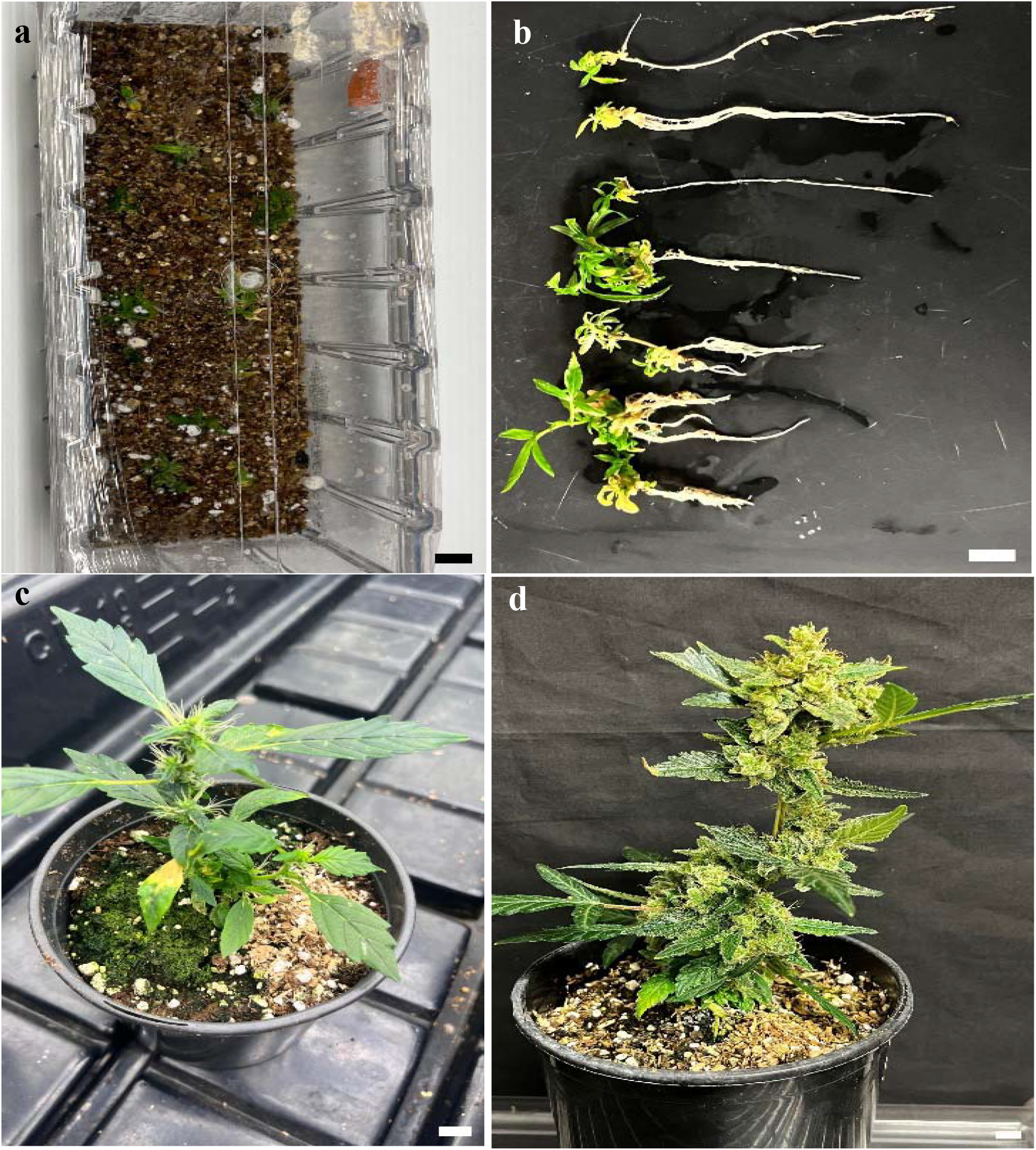
Acclimatization of reverted day-neutral plants a*) In vitro* plant in a vessel containing liquid DKW media + 4.92 μM IBA cultured in vermiculite, b) *In vitro* rooted plants after 21 days in medium, c) Plant in ex vitro condition after 30 days, d) Plant in ex vitro condition after 60 days. Scale bar: 1 cm.

### *In vitro* rooting media composition and acclimatization of the reverted shoots

Following the floral reversion process, some regenerated shoots were transferred into rooting medium. The rooting medium consisted of 30 grams of autoclaved vermiculate used as a substrate in We-V vessels (23 × 7 × 7 cm, We-V box; Magenta LLC, Chicago, IL) moistened with 120 mL of half-strength Murashige and Skoog modified Basal liquid medium (M530, PhytoTech Labs, Kansas, USA), 30 g/L sucrose, 1 mL/L plant preservative mixture (PPM), 6 g/L agar with 4.92 μM indole-3-butyric acid (IBA) according to (Ioannidis et al., 2022).

Lastly, a two-minute dip in autoclaved rooting powder containing 0.4% IBA (Plant Prod Stim Root #2 Brampton, Canada) was performed to enhance root development. Each vessel was placed under a photoperiod of 16/8 h (light/dark) at 25°C with a light intensity of approximately 30 μmol s−1 m−2(Fig 1a). After 21 days, shoot explants that had developed visible roots (Fig. 1b) were transferred to growth room conditions to acclimatize using standard protocols (Piunno et al., 2019). The plants were maintained under a long day photoperiod and developed to fully mature plants after 60 days of being transferred into the growth room (Fig. 1 c and d).

### *In vitro* rooting media composition and acclimatization of 2-year-old cultures

In addition to rooting shoots from the young florets used for the photoperiod experiments, this process was also replicated with florets from a day-neutral cultivar that had been maintained in tissue culture for over two years (subcultured approximately every three months). This 2-year-old culture had been maintained under the same *in vitro* conditions mentioned earlier in Magenta GA7 vessels containing floral reversion medium (DKW media with 1.0 μM meta-topolin hormone). The in vitro inflorescences were dissected into pairs of florets, and used to produce shoots that were subsequently rooted using the protocols mentioned in the previous sections (Ioannidis et al., 2022; Piunno et al., 2019) As with the other shoots, they were transferred to growth room conditions to acclimatize using standard protocols. The plants were maintained under a 16/8 (light/dark) photoperiod and developed into fully mature plants with fully developed seeds.

### Plant growth measurements

Each explant was monitored at the end of each week and floral reversion was indicated when the first shoot (minimum 3 mm) appeared from the explant, followed by leaf formation. All reverted floral explants were counted individually and divided by the total number of florets on each Petri dish to determine the average number of nodes and shoots per photoperiod treatment. To measure shoot length, reverted explants taken out of the medium were placed on an opaque colour sheet with a ruler beside them. After removing the initial florets and any callus, each plant was photographed and measured using ImageJ software (Version I.54f; National Institute of Mental Health, Bethesda, Maryland, USA). Node number and shoot number were carefully counted at the end of the experiment. The time of reversion for each explant was recorded weekly on day 7, day 14, day 21 and on the last day of the experiment on day 28^th^.

The floral reversion rate was obtained for all 25 Petri dishes by counting the number of reverted explants in one petri dish divided by the total number of explants per petri dish. This parameter was calculated weekly, on day 7, day 14, day 21 and day 28^th^ to determine the time of each floral explant undergoing floral reversion process.

### Experimental Design

This study was conducted in a customized shelving unit with a total of 15 chambers (3 × 5 rows) using a completely randomized design. Each chamber was equipped with LED lights that emit red, blue, green and yellow light. One side of each box had a small fan installed to provide ventilation and maintain temperature control. The back and front of each box were covered to prevent light leakage while allowing airflow. The entire unit was covered to avoid light leakage from the main grow chamber area to the experimental unit. One to a maximum of two Petri dishes were placed inside each box under LED lights (Fig.2s).

### Statistical analysis

This experiment is a completely randomized design (CRD) with five replicates. All data were analyzed using SAS Studio software (v9.4, SAS Institute Inc., Cary, NC, USA) Generalized linear mixed model function (PROC GLIMMIX with lognormal distribution) was employed for ANOVA, and LSMEANS statement(α = 0.05) was used for mean comparisons. Tukey–Kramer tests were employed to address multiple comparisons, and Microsoft Excel (Microsoft Corp., Redmond, WA, USA) was used for data visualization.

## Results

### Qualitative Observation of Shoot Proliferation under Different Photoperiods

Initial signs of shoot proliferation from floral explants were observed within the first week across all photoperiods to various degrees (Fig. 2). The explants under photoperiods 20/4, 16/8, and 18/6 showed small shoots with comparable sizes, indicating similar early-stage growth. However, the explants under 12/12 and continuous light (24/0) showed slightly smaller and less developed shoots.

**Fig 2.**
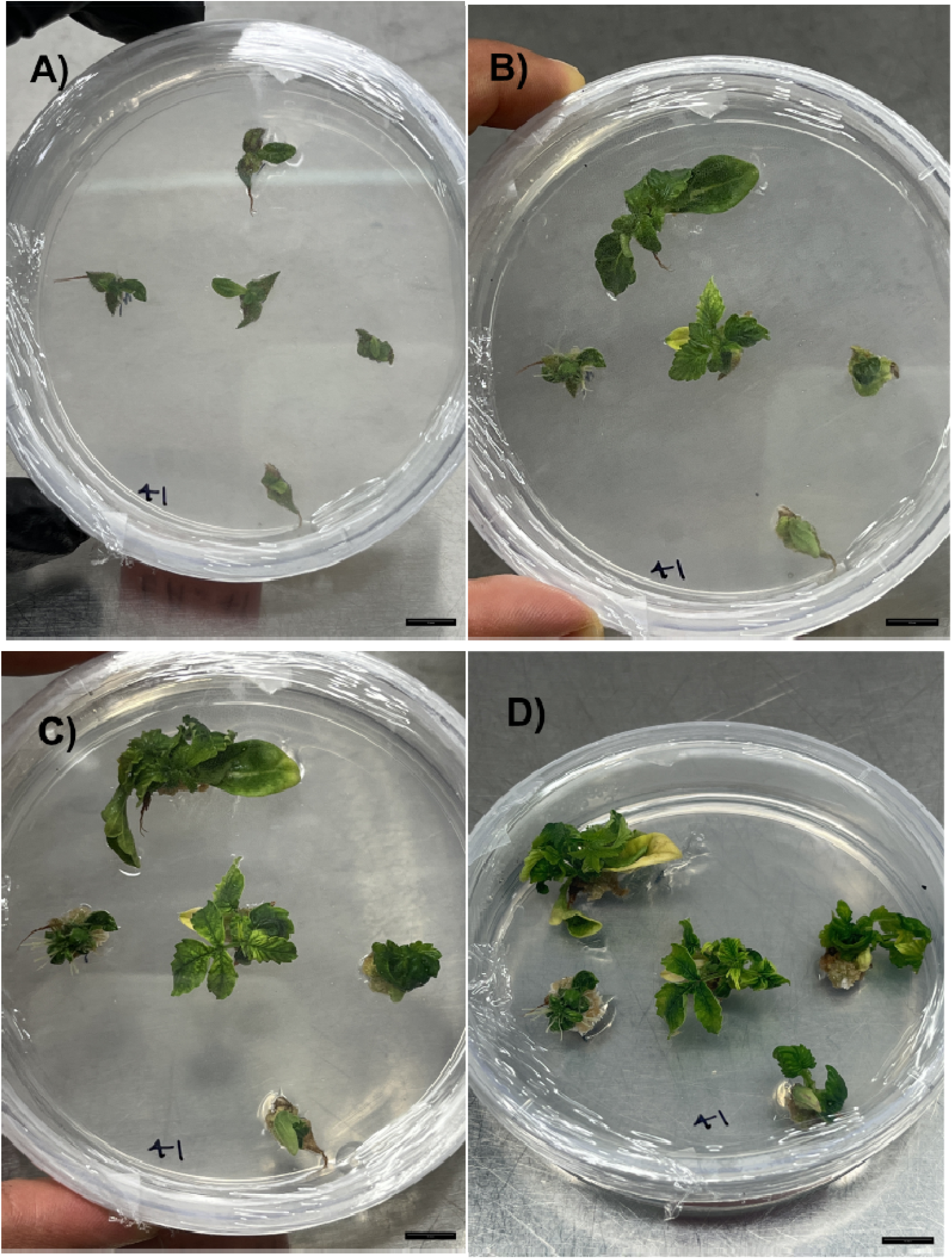
Representative photos of shoot proliferation from floral explants of day neutral *Cannabis sativa* cv Blue Auto Mazar x Auto Blueberry on DKW media enriched with 1 μM meta-topolin. Photos were taken on: A) day 7; B) day 14; C) day 21; D) day 28, under a 24/0 hr photoperiod. Scale bar: 1 cm.

By the second week, noticeable differences in shoot proliferation began. Explants under 20/4 and 16/8 exhibited more shoot growth with increased size and number, indicating these photoperiods promote early proliferation effectively. The explants under 12/12 showed moderate growth, while those under 18/6 and 24/0 demonstrated substantial increases in shoot size and number, suggesting that explants cultured under these photoperiods begin to catch up and these conditions promote more vigorous growth (Fig. 2).

In the third week, plants in all photoperiods showed further proliferation. Explants under 20/4, 16/8, and 18/6 exhibited increases in size, with well-developed structures. The explants under 12/12 showed moderate growth compared to the others, while those under 24/0 demonstrated substantial proliferation, with a notable increase in shoot number and size, suggesting that continuous light might accelerate growth after an initial slower phase (Fig. 2).

The differences became more pronounced by the end of the experiment in the fourth week. Explants under 20/4 and 16/8 displayed the most extensive and well-developed shoot proliferation, indicating these photoperiods are highly effective for sustained growth. The 18/6 photoperiod also resulted in well-developed shoots, comparable to 20/4 and 16/8. The 12/12 photoperiod, while showing consistent growth, resulted in less extensive proliferation compared to the others. Explants under continuous light (24/0) exhibited vigorous and abundant shoot development, suggesting that while initial growth might be slower, continuous light ultimately promotes extensive shoot proliferation (Fig. 2).

In addition, from the start of the third week to the fourth week, occasional friable non-organogenic callus formed in small amounts with yellow to brown colour. Callusing was mostly observed on the samples placed under longer photoperiods of 16/8, 18/6, 20/4 and 24/0 treatment. These calluses formed on the floret tissue surface where the shoots developed.

### Assessment of Floral Reversion Time and Photoperiod

The analysis of variance (ANOVA) result showed significant differences for the time of reversion (day) and its interactions in different photoperiods at the level of (*p <0*.*05*). The reversion rate on various days depended on the photoperiod. This result indicated that day 7 was significantly different (*p <0*.*05*) compared to days 14, 21 and 28 for photoperiods 16/8, 20/4 and 24/0, respectively. Meanwhile, during the first week of the experiment, explants under 12/12 hr of photoperiod showed the highest reversion rate with 52% reverted shoots followed by 18/6 hr with 48%, 16/8 hr of photoperiod with 36% and 20/4 and 24/0 hr with 20% and 12% respectively (Fig. 3).

**Fig 3.**
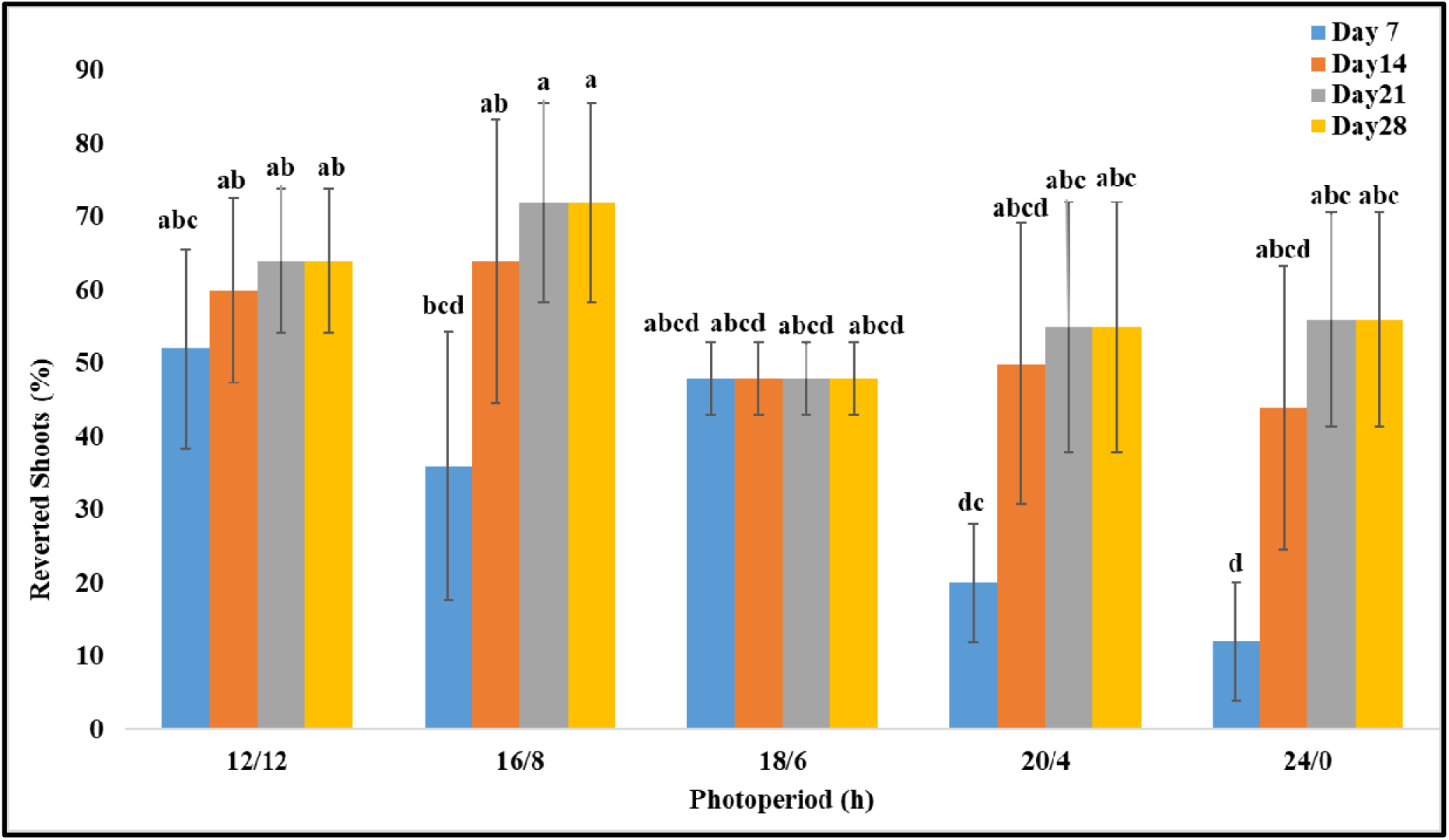
Percentage of shoot proliferation from floral explants at day 7, day 14, day 21 and day 28 of the experiment under five light treatments of 12/12, 16/8, 18/6, 20/4, and 24/0 hour photoperiods. Data are mean±SE (n = 5). Error bars indicate significant differences at *p* □*0*.*05* using the Tukey-Kramer multiple comparisons test.

After the second week, the highest revision rate was achieved under 16/8 hr of photoperiod with 64% of shoots reverted. The 12/12, 20/4 and 18/6 photoperiods showed 60%, 50% and 48% with the lowest reversion rate achieved under 24/0 hr of photoperiod with 44% of florets reverted after 14 days. The floral reversion rate was consistent on day 21 and day 28 for all photoperiod treatments. The highest amount was under the 16/8 photoperiod, reaching 72%. The 12/12 photoperiod also shows a high reversion rate, slightly lower than 16/8 with 64% of explants reverted. Other photoperiods result in lower reversion rates of 56% to 48% (Fig. 3).

### Evaluation of Shoot Length, Shoot Number and Node number

The analysis of variance (ANOVA) revealed there were no significant differences in terms of the photoperiod interaction among shoot length, shoot number and node number (*p <0*.*05*). While not significantly different, the shoot length of plants in the 20/4 hr photoperiod resulted in the longest average shoot length of 10.1 mm per explant (pair floret). Photoperiods 16/4 and 24/0 obtained slightly lower than the peak (10.1 mm) with 9.74 mm and 9.34 mm of shoot length per explant (Table 1). The shortest shoot lengths are observed under the 18/6 and 12/12 photoperiods, showing similar values of 6.72 mm and 5.78 mm respectively.

**Table 1.** The average shoot length(mm), shoot number and node number of reverted shoots obtained from five photoperiod treatments (12/12, 16/8, 18/6, 20/4, and 24/0 hour) measured on day 28. Each number represents the mean of 5 biological replicates +/-the standard error of the mean. None of the differences were statistically significant at *p* □*0*.*05*.

| Photoperiod (h) | Shoot length (mm) | Shoot number | Node number |
| --- | --- | --- | --- |
| | Mean ± SE | Mean | Mean ± SE |
| 12/12 | 5.78 ± 0.808 | 1 | 4 ± 0.316 |
| 16/8 | 9.74 ± 2.527 | 1 | 3 ± 0.316 |
| 18/6 | 6.72 ± 0.798 | 1 | 4.2 ± 0.2 |
| 20/4 | 10.1 ± 2.741 | 1 | 4.5 ± 0.447 |
| 24/0 | 9.34 ± 1.796 | 1 | 3.4 ± 0.6 |

For node number, similar to shoot length, plants grown under 20/4 hr developed the highest average of 4.5 nodes per shoot. Photoperiods 18/6 and 12/12 resulted slightly shorter with 4.2 and 4 nodes per explant. Fewer nodes (3.4) were observed under the continuous light treatment (24/0). Lastly, reverted explants under 16/8 photoperiod developed the fewest nodes on average per shoot (3). The results were similar for shoot number across all the photoperiod treatments where all the reverted explants developed one shoot per explant (Table 1). This result highlights that photoperiod had little effect on the floral reversion of day-neutral cannabis cv “BAM x AB”.

## Discussion

A majority of medicinal/recreational cannabis producers rely on clonal propagation to facilitate genetic and phenotypic uniformity. This is accomplished by maintaining mother plants in the vegetative stage under long photoperiods and propagating plants from vegetative tissues (Monthony et al., 2021). To reduce space, resources, and produce insect/disease-free propagules, nodal based micropropagation techniques have been developed (Hesami et al., 2023). However, this approach can only be achieved using photoperiodic cultivars and does not apply to day-neutral genotypes. In contrast, breeding and producing day-neutral cultivars relies on seed based propagation and breeding methods. Prior to this study, there were no published methods to propagate and maintain them in the vegetative stage as they shift into the flowering phase regardless of the photoperiod changes. The micropropagation method developed in this study represents a significant step toward large-scale clonal propagation for the production and controlled breeding of day-neutral cannabis.

Flower buds and other floral parts are effective explants for micropropagation in a variety of species (Ingvardsen et al., 2023). Floral reversion has been utilized for various plants as a simple and rapid way to regenerate plants using flowers as the starting material (Asker, 2016; Yongfeng et al., 2018). Date palms, *Arabidopsis*, Garlic, Lilies, and Switchgrass are only a few successful examples of floral reversion in plants (Asbe et al., 2015; Asker, 2016; Winiarczyk et al., 2018; Yongfeng et al., 2018; Zayed et al., 2016).

In many species, floral reversion occurs mostly through direct and indirect shoot organogenesis such as in *Mammillaria albicoma, Stevia rebaudiana, Agapanthus africanus, Allium giganteum*, and lilies (Ahmad et al., 2011; Asker, 2016; Supaibulwatana & Mii, 1997; Šušek et al., 2002; Wyka et al., 2006). While callus formation was observed to develop around some of the reverted explants in this study, previous work with photoperiodic cannabis identified that the reverted shoots developed from pre-existing meristems that subtend each floret and it is more likely that these are the source of shoots in this case (Monthony, et al., 2021). This observation is relevant since it is commonly believed that regeneration based micropropagation is more prone to somaclonal variation than systems that rely on shoot proliferation (Adamek et al., 2022).

Although floral reversion has been reported in cannabis, previous studies focussed exclusively on photoperiodic cultivars and no one has investigated floral reversion in day-neutral genotypes. The first successful floral reversion study on cannabis was done by (Piunno et al., 2019) using mature and immature greenhouse-sourced flowers. In this study, shoots were recovered from greenhouse flowers up until the day of harvest (Piunno et al., 2019). Following that study, Monthony et al., (2021) used *in vitro* florets and reported much higher reversion rates. This study investigated the effect of the size of floral explants, including single florets and pairs, and two cytokinins, mT and 6-benzylaminopurine (BAP) at various concentrations (Monthony, et al., 2021). After six to eight weeks in culture, compared to single florets, pairs of florets had a 2.5-3 times higher chance of reverting with 81% success in DKW medium enriched with 1 μM mT. This resulted in higher reversion rates and healthier explants; however, the timeframe was still lengthy and the process of dissecting florets was noted to be tedious and time-consuming (Monthony et al., 2021).

Our study builds on the previous floral reversion research by comparing the effect of various photoperiods on a day-neutral cultivar. Floral explants comprised of a pair of florets were cultured in DKW with 1 μmol mT and subjected to five photoperiods of 12/12, 16/8, 18/6, 20/4, and 24/0 hr (light/dark) treatments. Data analysis revealed a significant difference in terms of the interaction of photoperiod and the time of floral reversion. Photoperiods 16/8, 20/4, and 24/0 exhibited significant differences on the first week compared to weeks 2, 3 and 4 with reversion rates peaking at 72%, under 16/8 light treatments. Notably, the reversion rate remained consistent from week 3 to week 4, meaning the floral reversion process could be completed in three weeks. While there were some significant differences in the speed of reversion, our results highlight the lack of impact of photoperiod on final reversion rates, shoot elongation, and nodal development. This is not overly surprising given that day neutral cannabis plants do not use photoperiod for signalling, but this should be explored further using photoperiodic cultivars. Further, while increased daylength and daily light integral did not result in enhanced growth in this study, longer term experiments should be done.

Originally a period of 12 weeks was proposed for the floral reversion cycle for photoperiodic cultivars (Monthony et al., 2021). Although the floral reversion cycle for day-neutral genotypes was slightly different, a timeframe of 14 weeks is suggested for these cultivars (Fig. 4). This cycle started with the introduction of day-neutral seed to *in vitro* conditions. Flower induction of seed can take up to two months. The next step is the dissection of the floral explants and initiation of floral reversion which can take approximately four weeks. These shoots can then be grown in vitro to produce more florets to repeat the cycle and increase plant numbers or rooted and transferred ex vitro where they can be grown to maturity (Fig. 4). In addition to the experimental material used in the present study, a two-year-old *in vitro* culture of cv. “Blue Auto Mazar” (BAM) that had been maintained *in vitro* using floral reversion, was transferred to the growth room, acclimatized successfully, and grown into fully mature plants. The present study successfully demonstrated the first micropropagation protocol developed for day-neutral cannabis and will help facilitate large scale clonal propagation and breeding programs.

**Fig 4.**
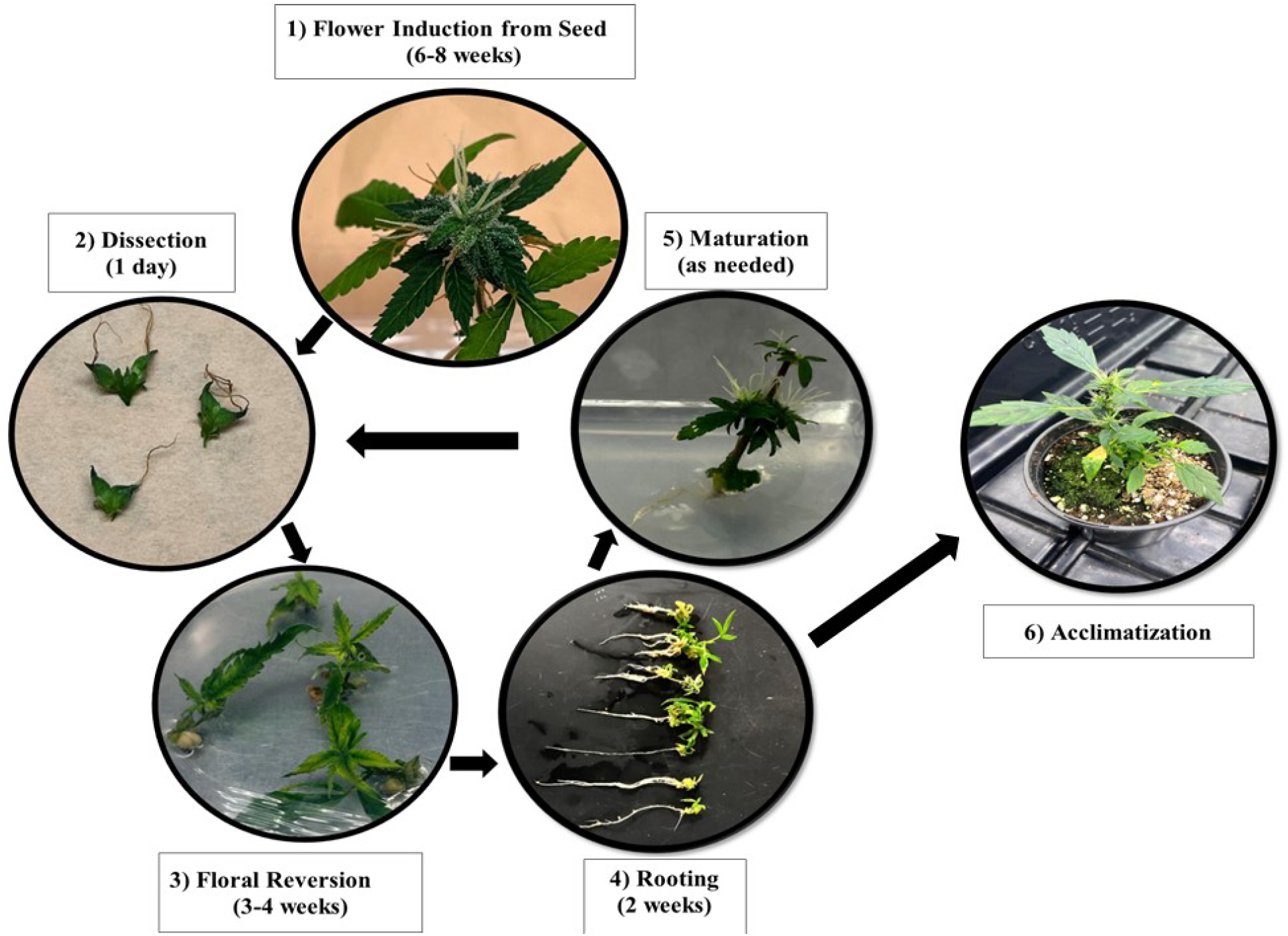
Presentation of *in vitro Cannabis sativa L*. Day-neutral Floral reversion cycle. 1) Flower induction from seeds: seeds germination and plant development in early stages of the reproductive stage for 6 to 8 weeks 2) Dissection: pairs of florets are dissected. 3) Floral reversion process: florets are cultured in DKW media with 1 μM meta-topolin hormone for 3 to 4 weeks.4) Rooting: Plants are rooted in vitro. 5) Maturation: shoots reverted from floral explants are cultured in MS rooting media and emerging flowers can be used again for floral reversion. 6) Acclimatization: rooted plants can be transferred to ex vitro conditions where they can complete their life cycle.

Future research studies could build upon the findings of this study to further enhance our understanding of floral reversion in *Cannabis sativa* L day-neutrals. One potential area of exploration is the use of multiple day-neutral cultivars to determine if the observed results are consistent across different genetic backgrounds, which would provide more generalizable conclusions. Future studies could also investigate the optimal number of florets numbers along with the best age of florets in various development stages. These follow-up studies would provide practical guidelines for efficient floral reversion and would further refine cannabis micropropagation techniques and implications for commercial cannabis cultivation and breeding programs.

## Conclusion

To the best of our knowledge, this study marks the first micropropagation system for day-neutral cultivars of cannabis. Breeding day neutrals is essentially restricted to population-level breeding in the absence of clonal propagation techniques needed for phenotyping and making F1 crosses as is done in photoperiodic cannabis. This protocol will facilitate targeted F1 crosses, elite selection inbreeding, and an overall increase in the pace of genetic gains. Importantly, reverted shoots, including shoots from a two-year-old culture of day-neutral cannabis, were successfully acclimatized in the greenhouse. While they appeared to enter reproductive growth quickly and produced small plants, they successfully completed their life cycle and could produce seeds. These findings underscore the potential of floral reversion as an efficient method for day-neutral cannabis propagation, providing a robust foundation for future research and commercial application in cannabis day-neutral cultivation.

## SUPPLEMENTARY INFORMATION

**Fig 1.**
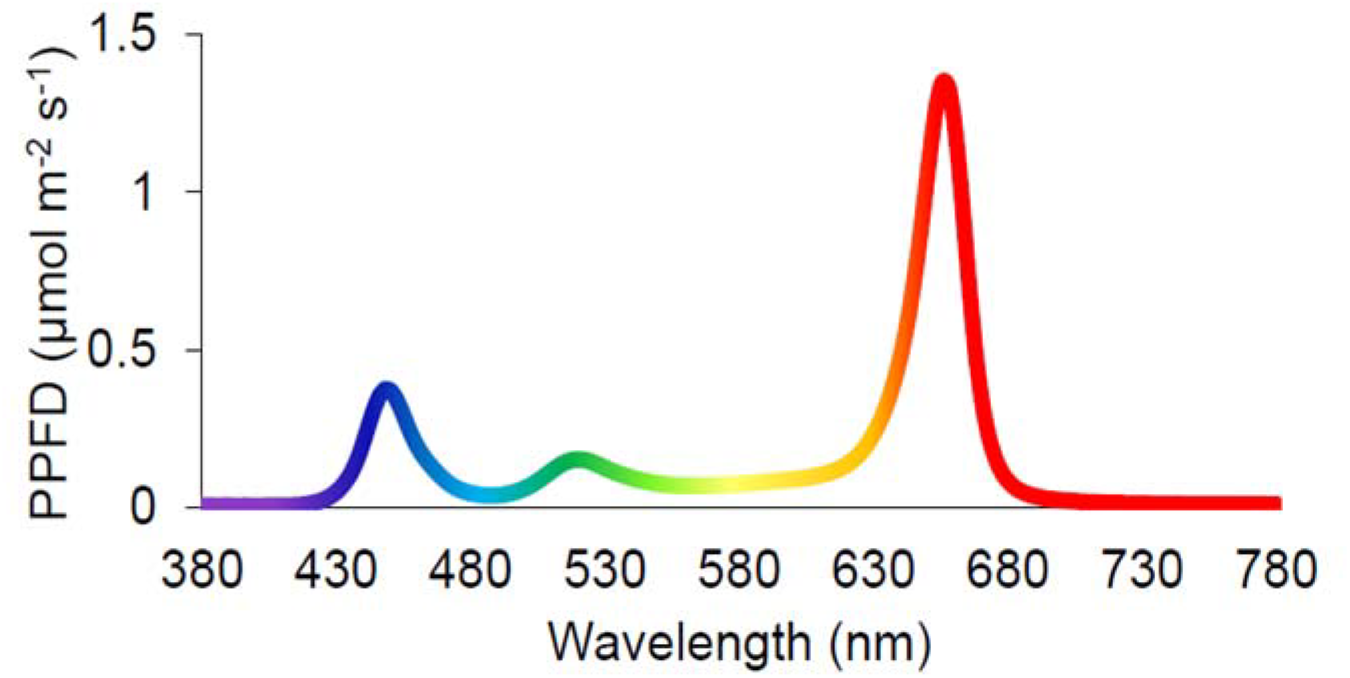
LI-COR LI-180 Spectrometer spectrum of LED illumination for reversing Cannabis sativa explants.

**Fig 2.**
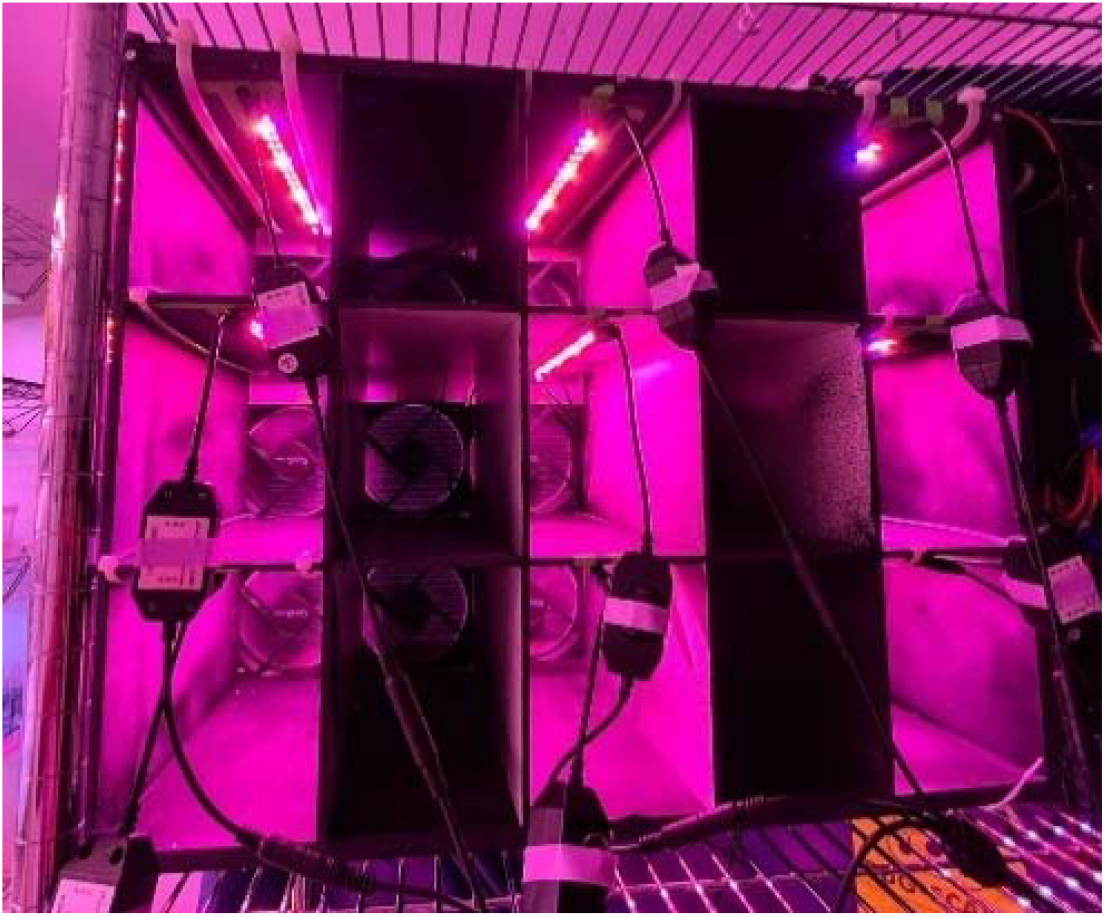
Representative photo of the experimental unit. Cables, fans and LED lights indicate a controlled environment. Each petri dish was placed inside of the boxes and the boxes were covered to prevent light leakage, yet allow ventilation to limit condensation.

## References

Adamek, K., Jones, A. M. P., & Torkamaneh, D. 2022. Accumulation of somatic mutations leads to genetic mosaicism in cannabis. Plant Genome, 15(1). 10.1002/tpg2.20169

Adhikary, D., Kulkarni, M., El-Mezawy, A., Mobini, S., Elhiti, M., Gjuric, R., Ray, A., Polowick, P., Slaski, J. J., Jones, M. P., & Bhowmik, P. 2021. Medical Cannabis and Industrial Hemp Tissue Culture: Present Status and Future Potential. In Frontiers in Plant Science (Vol. 12). Frontiers Media S.A. 10.3389/fpls.2021.627240

Ahmad, N., Fazal, H., Zamir, R., Khalil, S. A., & Abbasi, B. H. 2011. Callogenesis and Shoot Organogenesis from Flowers of Stevia rebaudiana (Bert.). Sugar Tech, 13(2), 174–177. 10.1007/s12355-011-0083-3

Ahrens, A., Llewellyn, D., & Zheng, Y. 2023. Is Twelve Hours Really the Optimum Photoperiod for Promoting Flowering in Indoor-Grown Cultivars of Cannabis sativa? Plants, 12(14), 2605. 10.3390/plants12142605

Anna K. Jägera, A.H. bJ. van S.a. 1995. Screening of Zulu medicinal plants for prostaglandin-synthesis inhibitors. Journal of Ethnopharmacology.

Asbe, A., Matsushita, S. C., Gordon, S., Kirkpatrick, H. E., & Madlung, A. 2015. Floral reversion in Arabidopsis suecica is correlated with the onset of flowering and meristem transitioning. PLoS ONE, 10(5). 10.1371/journal.pone.0127897

Asker, H. M. 2016. In vitro direct organogenesis in response to floral reversion in lily. African Journal of Biotechnology, 15(44), 2497–2506. 10.5897/ajb2016.15639

Babaei, M., Ajdanian, L., & Asgari Lajayer, B. 2022. Morphological and phytochemical changes of Cannabis sativa L. affected by light spectra. In New and Future Developments in Microbial Biotechnology and Bioengineering (pp. 119–133). Elsevier. 10.1016/b978-0-323-85581-5.00020-3

Bonini, S. A., Premoli, M., Tambaro, S., Kumar, A., Maccarinelli, G., Memo, M., & Mastinu, A. 2018. Cannabis sativa: A comprehensive ethnopharmacological review of a medicinal plant with a long history. In Journal of Ethnopharmacology (Vol. 227, pp. 300–315). Elsevier Ireland Ltd. 10.1016/j.jep.2018.09.004

Caplan, D., Stemeroff, J., Dixon, M., & Zheng, Y. 2018. Vegetative propagation of cannabis by stem cuttings: effects of leaf number, cutting position, rooting hormone, and leaf tip removal. 10.1139/cjps-2018-0038

Hammond, C. T., & Mahlberg, P. G. 1977. Morphogenesis of Capitate Glandular Hairs of Cannabis sativa (Cannabaceae). In Source: American Journal of Botany (Vol. 64, Issue 8). https://about.jstor.org/terms

Handbook of Cannabis Production in Controlled Environments. (n.d.).

Hesami, M., Adamek, K., Pepe, M., & Jones, A. M. P. 2023. Effect of Explant Source on Phenotypic Changes of In Vitro Grown Cannabis Plantlets over Multiple Subcultures. Biology, 12(3). 10.3390/biology12030443

Hesami, M., Pepe, M., Baiton, A., & Jones, A. M. P. 2023. Current status and future prospects in cannabinoid production through in vitro culture and synthetic biology. In Biotechnology Advances (Vol. 62). Elsevier Inc. 10.1016/j.biotechadv.2022.108074

Hesami, M., Pepe, M., Monthony, A. S., Baiton, A., & Phineas Jones, A. M. 2021. Modeling and optimizing in vitro seed germination of industrial hemp (Cannabis sativa L.). Industrial Crops and Products, 170. 10.1016/j.indcrop.2021.113753

Ingvardsen, C. R., & Brinch-Pedersen, H. 2023. Challenges and potentials of new breeding techniques in Cannabis sativa. In Frontiers in Plant Science (Vol. 14). Frontiers Media S.A. 10.3389/fpls.2023.1154332

Ioannidis, K., Tomprou, I., & Mitsis, V. 2022. An Alternative In Vitro Propagation Protocol of Cannabis sativa L. (Cannabaceae) Presenting Efficient Rooting, for Commercial Production. Plants, 11(10). 10.3390/plants11101333

Kalant, O. J. 1972. Report of the Indian Hemp Drugs Commission, 1893-94: A Critical Review. International Journal of the Addictions, 7(1), 77–96. 10.3109/10826087209026763

Livingston, S. J., Quilichini, T. D., Booth, J. K., Wong, D. C. J., Rensing, K. H., Laflamme-Yonkman, J., Castellarin, S. D., Bohlmann, J., Page, J. E., & Samuels, A. L. (2020). Cannabis glandular trichomes alter morphology and metabolite content during flower maturation. Plant Journal, 101(1), 37–56. 10.1111/TPJ.14516

Moher, M., Jones, M., & Zheng, Y. 2021. Photoperiodic response of in vitro cannabis sativa plants. HortScience, 56(1), 108–113. 10.21273/HORTSCI15452-20

Monthony, A. S., Bagheri S, Zheng Y, & Jones A. M. P. 2021. Flower Power: Floral reversion as a viable alternative to nodal micropropagation in Cannabis sativa. 10.1101/2020.10.30.360982

Monthony, A. S., Page, S. R., Hesami, M., Maxwell, A., & Jones, P. 2021. plants The Past, Present and Future of Cannabis sativa Tissue Culture. 10.3390/plants

Piunno, K. F., Golenia, G., Boudko, E. A., Downey, C., & Maxwell, A. (2019a). Regeneration of shoots from immature and mature inflorescences of cannabis sativa. Canadian Journal of Plant Science, 99(4), 556–559. 10.1139/cjps-2018-0308

Poluboyarova, T. V., Novikova, T. I., Vinogradova, G. Yu., & Andronova, E. V. 2014. Morpho-Histological Analysis of Direct Shoot Organogenesis Induced in Flower Buds Cultures of *Allium altissimum* American Journal of Plant Sciences, 05(13), 2015–2022. 10.4236/ajps.2014.513216

Spitzer-Rimon, B., Duchin, S., Bernstein, N., & Kamenetsky, R. 2019. Architecture and florogenesis in female Cannabis sativa plants. Frontiers in Plant Science, 10. 10.3389/fpls.2019.00350

Spitzer-Rimon, B., Shafran-Tomer, H., Gottlieb, G. H., Doron-Faigenboim, A., Zemach, H., Kamenetsky-Goldstein, R., & Flaishman, M. 2022. Non-photoperiodic transition of female cannabis seedlings from juvenile to adult reproductive stage. Plant Reproduction, 35(4), 265–277. 10.1007/s00497-022-00449-0

Supaibulwatana, K., & Mii, M. 1997. Organogenesis and Somatic Embryogenesis from Young Flower Buds of Agapanthus africanus Hoffmanns. In Original Papers Plant Biotechnology (Vol. 14, Issue 1).

Šušek, A., Javornik, B., & Bohanec, B. 2002. Factors affecting direct organogenesis from flower explants of Allium giganteum. In Plant Cell, Tissue and Organ Culture (Vol. 68).

Winiarczyk, K., Marciniec, R., & Tchórzewska, D. 2018. Phenomenon of floral reversion in bolting garlic (Allium sativum L.). Acta Scientiarum Polonorum, Hortorum Cultus, 17(2), 123–134. 10.24326/asphc.2018.2.11

Wyka, T. P., Hamerska, M., & Wróblewska, M. 2006. Organogenesis of vegetative shoots from in vitro cultured flower buds of Mammillaria albicoma (Cactaceae). Plant Cell, Tissue and Organ Culture, 87(1), 27–32. 10.1007/s11240-006-9128-9

Yongfeng, W., Aiquan, Z., Fengli, S., Mao, L., Kaijie, X., Chao, Z., Shudong, L., & Yajun, X. 2018. Using transcriptome analysis to identify genes involved in switchgrass flower reversion. Frontiers in Plant Science, 871. 10.3389/fpls.2018.01805

Zayed, E. M. M., Zein El Din, A.F.M., Manaf, H. H., & Abdelbar, O. H. 2016. Floral reversion of mature inflorescence of date palm in vitro. Annals of Agricultural Sciences, 61(1), 125–133. 10.1016/j.aoas.2016.01.003

